# Carboxydovores from the Pseudomonadota colonise volcanic soils during succession

**DOI:** 10.1101/2023.11.12.566731

**Authors:** Robin A. Dawson, Patricia Aguila, Gary M. King, Marcela Hernández

## Abstract

Carbon monoxide (CO) degrading microorganisms are present in volcanic deposits throughout succession, with vegetation and soil influencing the communities present. The carboxydovores are a subset of CO degraders that use CO only as an energy source, raising the question of how the physiological and metabolic features of the carboxydovores can make these bacteria more competitive in harsh volcanic ecosystems. An enrichment strategy was modified, which enabled the isolation of two carboxydovore representatives from genera that were abundant in the native soils, *Cupriavidus* sp. CV2^T^ (92.3% ANI vs. *Cupriavidus basilensis* DSM 11853) and a putative strain of *Paraburkholderia terrae* (*Pb. terrae* COX) (96.42% ANI vs. *Pb. terrae* KU-64^T^). These isolates oxidise CO across a very broad range of concentrations, and genome sequence analysis indicated that they use form-I carbon monoxide dehydrogenase (CODH) to do so. *Cupriavidus* sp. CV2^T^ and *Pb. terrae* COX each oxidised CO specifically at stationary phase, but the conditions for induction of CODH expression were distinct. *Cupriavidus* sp. CV2^T^ expressed CODH only in the presence of CO, while *Pb. terrae* COX expressed CODH regardless of the presence of CO. Based on metabolic and phylogenetic analyses, *Cupriavidus* sp. CV2^T^ is recommended as a novel species within the genus *Cupriavidus*. Therefore, we propose the name *Cupriavidus ulmosensis* sp. nov. for the type strain CV2^T^ (= NCIMB 15506^T^, = CECT 30956^T^). This study provides valuable insights into the physiology and metabolism of carboxydovores, which colonise volcanic ecosystems during succession.

**Importance:** Volcanic ecosystems harbour many bacteria that contribute to the environmentally important process of carbon monoxide (CO) oxidation. We demonstrate a modified method for isolating bacteria, which consume CO at very low concentrations as a supplementary energy source (carboxydovory), leading to the isolation of two novel strains (*Cupriavidus* sp. CV2^T^ and *Paraburkholderia terrae* COX) from volcanic strata that formed in 1917 and 2015, respectively. The conditions under which CO consumption occurs were investigated; each strain consumed CO during stationary phase, but *Pb. terrae* COX consumed CO regardless of the prior growth conditions while *Cupriavidus* sp. CV2 was more controlled. *Cupriavidus* sp. CV2 is a type strain of a new species, *Cupriavidus ulmosensis* str. CV2, which demonstrates relatively high tolerance for CO. These strains provide the basis for further study of the physiology, metabolism, and genetics of CO oxidation by carboxydovores, and will help us to understand how bacteria colonise harsh volcanic ecosystems.

## INTRODUCTION

Carbon monoxide (CO) is a toxic gas, which is emitted to the atmosphere in large quantities each year (∼2,600 Tg yr^-1^ emitted) (Khalil and Rasmussen, 1990), causing significant changes to atmospheric chemistry and climate. Levels of CO in non-urban environments typically range from 60-300 ppbv (King, 2003b), but anthropogenic sources can increase urban CO concentrations into the low ppmv range (Wright *et al*., 1975; Saadi *et al*., 2021), causing issues with air quality. Up to 2,100 Tg CO yr^-1^ is removed abiotically from the atmosphere through reactions with hydroxyl radicals, hindering the subsequent removal of polluting compounds (e.g. methane) (Crutzen and Gidel, 1983) and exacerbating the effects of global warming.

Soils act as a significant sink for CO due to microbial activity (Conrad and Seiler, 1980; Bartholomew and Alexander, 1981), accounting for removal of ∼250-300 Tg yr^-1^ (King and Weber, 2007; Cordero *et al*., 2019). Both aerobic and anaerobic microorganisms are capable of CO oxidation due to the use of a Mo-Cu or Ni-Fe carbon monoxide dehydrogenases (CODH), respectively (King, 2003b; Depoy *et al*., 2021), although limited evidence is available to suggest that anaerobic CO degraders play a role in atmospheric CO consumption. CO degraders colonise diverse ecological niches, and isolates have been retrieved from terrestrial, aquatic and even clinical samples (King and Weber, 2007; Stott *et al*., 2008; King and King, 2014). CO degraders can be broadly classified into two groups according to the specific usage of CO; carboxydotrophs and carboxydovores. Carboxydotrophs can grow using CO as the sole source of carbon and energy, with many isolates able to use high concentrations of CO (e.g. 10% (v/v) (King and Weber, 2007), and can tolerate up to 90% (v/v) headspace CO (Cypionka and Meyer, 1982; Mörsdorf *et al*., 1992), although some carboxydotrophs perform better with lower CO concentrations (e.g. 3% v/v) (Cypionka and Meyer, 1982). It is unclear whether carboxydotrophs are able to oxidise atmospheric concentrations of CO, although early studies suggested that they cannot (Conrad, 1996). CODH catalyses the oxidation of CO to CO_2_, meaning that carboxydotrophs can utilise a pathway for carbon fixation such as the Calvin-Benson-Basham cycle to metabolise the generated CO_2_ while also gaining ATP and NAD(P)H + H^+^ (Siebert *et al*., 2020). Conversely, while carboxydovores may oxidise higher concentrations of CO, they can also oxidise atmospheric levels, but only benefit from this reaction by acquiring energy in the form of electrons (King, 2003b; Cordero *et al*., 2019). Carboxydovores often lack the genes required for fixation of CO_2_, but some reports indicate that these CO degraders are actually inhibited by the high concentrations of CO required by carboxydotrophs for growth (Hardy and King, 2001; Weber and King, 2007).

The physiology, biochemistry and genetics of carboxydotrophs have been studied in detail, with many studies focussing on the model bacterium *Oligotropha carboxidovorans* OM5^T^ (Schubel *et al*., 1995; Nam *et al*., 2003; Wilcoxen and Hille, 2013; Siebert *et al*., 2020). However, relatively few publications are available that focus on bacterial carboxydovores in similar detail (King, 2003b; King and King, 2014; Cordero *et al*., 2019; Islam *et al*., 2019). Recent studies have begun to demonstrate that carboxydovores may be abundant and environmentally important bacteria (Quiza *et al*., 2014), especially as this group are able to oxidise atmospheric concentrations of CO (Islam *et al*., 2019), but the contribution of carboxydovores to CO uptake *in situ* remains to be established. We require a greater diversity of cultured representatives to fully appreciate the contribution of carboxydovores to the consumption of atmospheric CO. In this study, a targeted enrichment strategy was modified and employed on volcanic soils, leading to the isolation of two novel carboxydovores from the Pseudomonadota (formerly Proteobacteria). According to 16S rRNA amplicon sequence analysis, these isolates are representatives of abundant taxa within the native volcanic deposits. Physiological, biochemical, and genomic analyses revealed the metabolic flexibility of these bacteria, as well as their distinctive use of CO in response to carbon limitation.

## RESULTS AND DISCUSSION

### Targeted enrichment and isolation of carboxydovores

In this study, an enrichment strategy was modified (from Weber and King, 2017) to isolate carboxydovores from volcanic strata formed after eruptions in 1917 (regenerated soil) and 2015 (mostly rock). The enrichment conditions were designed to promote mixotrophy, using 0.5 mM pyruvate to facilitate growth and 100 ppmv CO to provide supplementary energy for CO degrading bacteria. 100 ppmv CO was deemed to be too low to enrich carboxydotrophs, as very high concentrations of CO (e.g. 10-20% (v/v)) are used by these bacteria for growth (King, 2003b). Although heterotrophic bacteria would have been able to utilise the pyruvate, this enrichment strategy relied on the ability of carboxydovores to remain competitive by acquiring supplementary energy through CO oxidation, as suggested previously (Weber and King, 2010a). 0.5 mM pyruvate was selected as higher concentrations of organic carbon may inhibit CO consumption (Weber and King, 2007, 2012), but concentrations of pyruvate around 0.4 mM still permitted CO oxidation by members of the Burkholderiales (Weber and King, 2012). Similarly, *Labrenzia aggregata* consumed CO when grown with glucose concentrations ≤0.5 mM, suggesting that low concentrations of heterotrophic carbon would be suitable for a range of CO degraders (Weber and King, 2007). The enrichment conditions permitted CO oxidation by the microbial communities in Calbuco volcanic soils and sediments, as ∼100 ppmv CO was consumed after 48 and 72 hours by the 1917 and 2015 stratifications, respectively (data not shown). CO was consumed by slurry enrichments and sub-cultured enrichments over the course of 700-hour incubations.

The carboxydovores, *Cupriavidus* sp. CV2^T^ and *Pb. terrae* COX, were isolated from 1917 and 2015 volcanic stratifications, respectively. *Cupriavidus* spp. have previously been isolated from volcanic ecosystems due to their ability to oxidise H_2_, but they have not been isolated as carboxydovores (Sato *et al*., 2006). Previously, *Cupriavidus necator* was noted for containing a form-II CODH, but this form is not certain to oxidise CO as a physiological substrate (King and Weber, 2007). *C. necator* H16 grows autotrophically with CO_2_ and H_2_ and, although it was able to tolerate high CO concentrations, it could not oxidise CO to CO_2_ (Wickham-Smith *et al*., 2023). Conversely, CO oxidation by *Burkholderia* and *Paraburkholderia* species from volcanic soils has been well established (Weber and King, 2010b, 2017), although *Paraburkholderia terrae* has not been previously identified as a CO degrader.

*Cupriavidus* sp. CV2^T^ and *Pb. terrae* COX are both members of the Pseudomonadota, a group that was previously noted for being highly abundant in vegetated volcanic deposits (Dunfield and King, 2004; Weber and King, 2010a; Hernández *et al*., 2020a). The Pseudomonadota may be important to environmental CO cycling as, in vegetated volcanic soils, an increase in Pseudomonadota *coxL* sequences correlated with higher CO uptake activity (Weber and King, 2010a). The 2015 samples were almost entirely rock, with only traces of newly-formed soil present, likely necessitating the use of reduced gases such as CO_2_, CO and H_2_ by the resident microbial community (Fierer *et al*., 2010; Hernández *et al*., 2020b). Recently formed soil on volcanic sediments, such as the 1917 soils, provides a wealth of organic carbon to the resident microbial community, coinciding with increased Pseudomonadota sequences (Weber and King, 2010a). Weber & King (2010a) posited that CO oxidisers were able to compete with other heterotrophs for organic carbon during ecosystem development, and CO is supplied to these communities via plant roots (King and Crosby, 2002). However, vegetated Hawaiian volcanic deposits were net emitters of atmospheric CO (King and Weber, 2008) in spite of the high potential CO uptake potential in these soils, possibly due to abiotic CO production associated with high organic matter concentrations. Grasses colonised the vertical surface of the Calbuco stratification (Figure S1), potentially providing a source of CO to the resident microbial community. However, it must be noted that *in situ* CO fluxes were not measured in this study.

### Relative abundance of the Pseudomonadota in recent volcanic deposits

The bacterial communities that inhabited the 2015 and 1917 volcanic deposits were largely dominated by the Pseudomonadota (26.08% and 28.99% average relative abundance (RA), respectively), Actinomycetota (29.09% and 5.21% RA, respectively) and Acidobacteriota (10.79% and 26.02% RA, respectively) (Figure 1A). Other groups of note included the Chloroflexota, previously noted for its abundance in recent volcanic deposits (Weber and King, 2010a; Hernández *et al*., 2020a; Dragone *et al*., 2023) and its potential for CO oxidation (King and King, 2014; Islam *et al*., 2019; Hernández *et al*., 2020b). The Chloroflexota increased from a RA of 0.96% in the 2015 deposit to 8.01% in the 1917 deposit, indicating that this group was less suited to colonising the very earliest volcanic deposit (2015), but was more competitive during soil succession. The Acidobacteriota have been detected in relatively high abundance in vegetated and unvegetated volcanic deposits from Llaima and Kilauea volcanoes (Weber and King, 2010a; Hernández *et al*., 2020a). The Pseudomonadota were the most abundant group in recent deposits from the Krafla volcanic field (Iceland) (Byloos *et al*., 2018) and in deposits from Miyake-Jima (Japan) (Guo *et al*., 2014). Members of the Pseudomonadota were also abundant in deposits from the older/revegetated Llaima and Kilauea volcanoes (Weber and King, 2010a; Hernández *et al*., 2020a). This contrasts with the relative abundance of the Pseudomonadota observed in the unvegetated 2015 deposit in Calbuco (Figure 1A). The reason for the differing abundance of the Pseudomonadota in young volcanic deposits is unclear, although it was previously suggested that the abundance of this group increased with increasing availability of organic carbon (Weber and King, 2010a). As *Cupriavidus* sp. CV2^T^ and *Pb. terrae* COX are members of the Pseudomonadota, the abundance of this group was studied in greater detail to determine the potential importance of these strains *in situ*.

**Figure 1.**
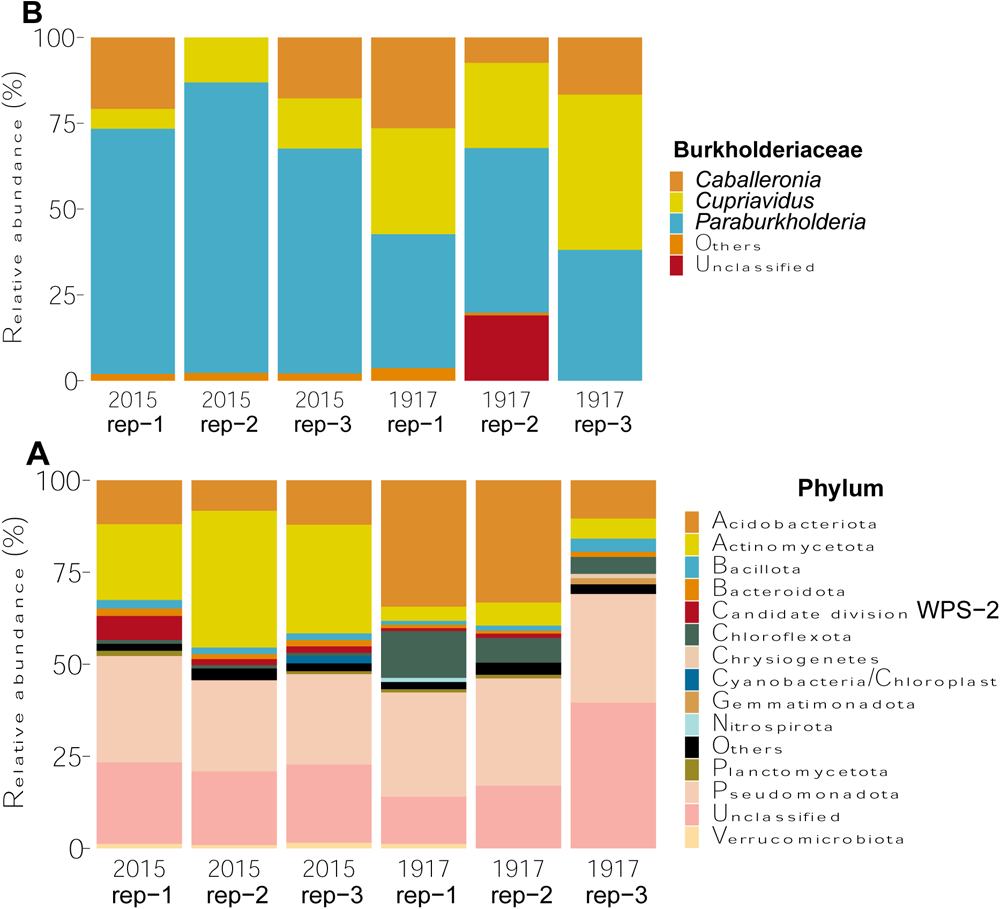
Relative abundance of 16S rRNA gene sequences in Calbuco stratifications originating from eruptions in 2015 and 1917. A) Relative abundance of 16S rRNA divided by phylum. B) Relative abundance of members of the Burkholderiaceae. Where the abundance of unclassified was lower than 0.8%, it was grouped into “Others”.

Alphaproteobacteria dominated both the 2015 and 1917 volcanic deposits (32.97% and 34.99% RA, respectively), followed by Betaproteobacteria (12.22% and 18.64% RA, respectively) (data not shown). Within the Betaproteobacteria, Burkholderiaceae was dominated by the genus *Paraburkholderia* in the 2015 volcanic deposit (73.81% average RA), with a decrease in abundance in the 1917 deposit (41.67% RA) (Figure 1B). These data indicate that *Pb. terrae* is a member of a well-represented group bacteria *in situ*. The genera *Burkholderia* and *Paraburkholderia* were previously suggested to be important CO oxidisers within the Pseudomonadota (Weber and King, 2010a, 2017), and this study provides valuable evidence that CO-oxidising members of this group can be present in young, unvegetated volcanic deposits. *Cupriavidus* spp. were less abundant in the 2015 volcanic deposit (11.17% RA), but increased to 33.64% RA in the 1917 volcanic deposit (Figure 1B), indicating that *Cupriavidus* sp. CV2^T^ is relatively well-represented in the older soil. These data are expected, as *Cupriavidus*-related bacteria, many of which oxidised CO_2_ and H_2_, comprised 25% of culturable bacteria from volcanic deposits from Mt. Pinatubo (the Philippines) (Ogiwara *et al*., 1999; Sato *et al*., 2004, 2006).

### Genome sequence analysis of novel carboxydovores

Strains *Cupriavidus* sp. CV2^T^ and *Pb. terrae* COX have similarity to *Cupriavidus basilensis* strain DSM 11853^T^ (99.33% nucleotide ID with 16S rRNA, BioSample ID SAMN12697569, Bioproject PRJNA563568) and *Paraburkholderia terrae* strain KU-64^T^ (NBRC 100964, 99.73%, Biosample ID SAMD00391168, Bioproject PRJNA33175). The genome of *Cupriavidus* sp. CV2^T^ has a completeness of 99.5 % with 2.3% contamination. The genome of *Paraburkholderia terrae* COX has a completeness of 99.7% with 2.4% contamination. Additional genome information is listed in Table S1. It is interesting that *Paraburkholderia terrae* strain COX was isolated from a recently deposited volcanic soil (lava 2015), as Weber and King (2010b) did not detect *Burkholderia*-like *coxL* sequences in bare Hawaiian samples.

In order to determine whether either carboxydovore was a member of a new species, genome-based taxonomic analysis was conducted using TYGS (Figures S2, S3), Average Nucleotide Identity (ANI) and digital DNA-DNA hybrisisation (dDDH) as well as autoMLST. *Cupriavidus* sp. CV2^T^ showed an ANI of 92.3% and a dDDH of 47.5% to their closest type strain *Cupriavidus basilensis* strain DSM 11853 (data not shown). *Pb. terrae* COX showed an ANI of 96.42% and a dDDH of 71.1% to their closest type strain *Paraburkholderia terrae* strain KU-64 (NBRC 100964, data not shown). *Cupriavidus* sp. CV2^T^ was determined to be a member of a new species, one that was closely related to the type strain *Cupriavidus basilensis* strain DSM 11853. *Pb. terrae* COX, however, was confirmed to be a *Paraburkholderia terrae*. CO-oxidising *Paraburkholderia* spp. have previously been isolated from soils from Kilauea volcano (1959 deposit, vegetated) (Weber and King, 2017).

The genome of *Cupriavidus* sp. CV2^T^ is over 10 Mbp with a GC content of 64.8% (Table S1) compared with 8.9 Mbp and a GC content of 65.0% for the type strain *Cupriavidus basilensis* DSM 11853 (Salvà-Serra *et al*., 2021). The genome of *Pb. terrae* COX is over 10 Mbp with a GC content of 62.0%, similar to the type strain *Paraburkholderia terrae* strain KU-64 (NBRC 100964), which has a genome measuring 9.9 Mbp with a GC content of 61.5%. The genomes of *Cupriavidus* sp. CV2^T^ and *Pb. terrae* COX contain a large number of coding sequences (10,380 and 10,390, respectively), and subsystems analysis indicated that many of these genes were devoted to metabolism (Table S1). *Cupriavidus* sp. CV2^T^ contains more genes with predicted roles involved in membrane transport and fatty acid metabolism, while *Pb. terrae* COX encoded more genes with predicted roles in iron acquisition and metabolism and carbohydrate metabolism (Table S1). The environmental source of each strain provides valuable context when considering their metabolic potential; *Cupriavidus* sp. CV2^T^ was likely to encounter more organic material in its natural environment, while *Pb. terrae* COX would have to employ a versatile metabolism in order to take the fullest advantage of transient nutrient sources and trace gases. CO flux measurements were not taken at Calbuco volcano, but previous studies have demonstrated that significant quantities of CO are produced by plant roots and through abiotic processes (King and Crosby, 2002; King and Weber, 2008).

Consistent with the isolation of *Cupriavidus* sp. CV2^T^ and *Pb. terrae* COX during enrichment with CO, visually identical gene clusters were identified on each genome that were highly likely to encode a form-I CODH (*coxMSL*) (Figure 2). *coxMSL* encodes the medium (FAD-binding), small (2 x [2Fe-2S]) and large (Mo-Cu) subunits of carbon monoxide dehydrogenase, respectively, with CoxL containing the active site and critical molybdopterin cystosine dinucleotide cofactor (Johnson *et al*., 1990; Meyer *et al*., 1993). The form of *coxL* was determined by identifying the active site motif AYXCSFR, characteristic of form-I CODH (Dunfield and King, 2004), during alignment of the translated amino acid sequences with known form-I CoxL sequences. CoxL amino acid sequences for *Cupriavidus* sp. CV2^T^ and *Pb. terrae* COX were most closely related to other Pseudomonadota CoxL sequences (Figure 3). tBLASTn analysis of the *coxL* gene from *Cupriavidus* sp. CV2^T^ against the genomes of *Cupriavidus* spp. type strains yielded varying results; *Cupriavidus basilensis* DSM 11853^T^ had no *coxL* genes or *coxMSL* gene clusters, while *Cupriavidus necator* N-1^T^ and *Cupriavidus oxalaticus* strain Ox1^T^ each encoded form-II CODH gene clusters (*coxSLM*), indicating that CO oxidation using form-I CODH may be unusual in the genus *Cupriavidus*.

**Figure 2.**
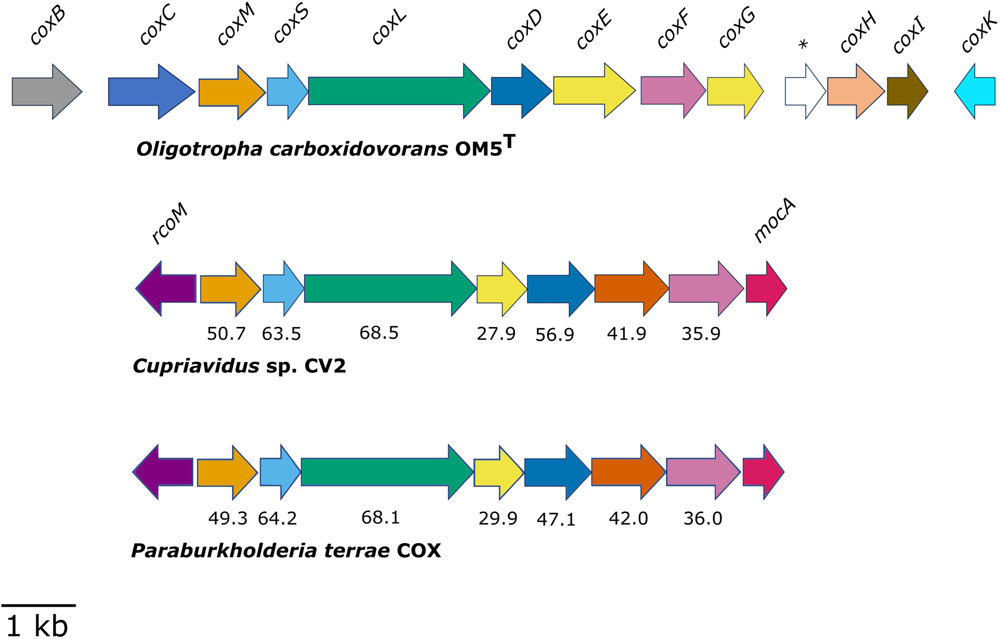
Form-I CODH-encoding gene clusters from *Cupriavidus* sp. CV2^T^ and *Paraburkholderia terrae* COX. Translated amino acid identities (%) were calculated using BLASTp against homologous query sequences from the model type strain carboxydotroph *Oligotropha carboxidovorans* OM5. Asterisk (*) indicates a hypothetical gene.

**Figure 3.**
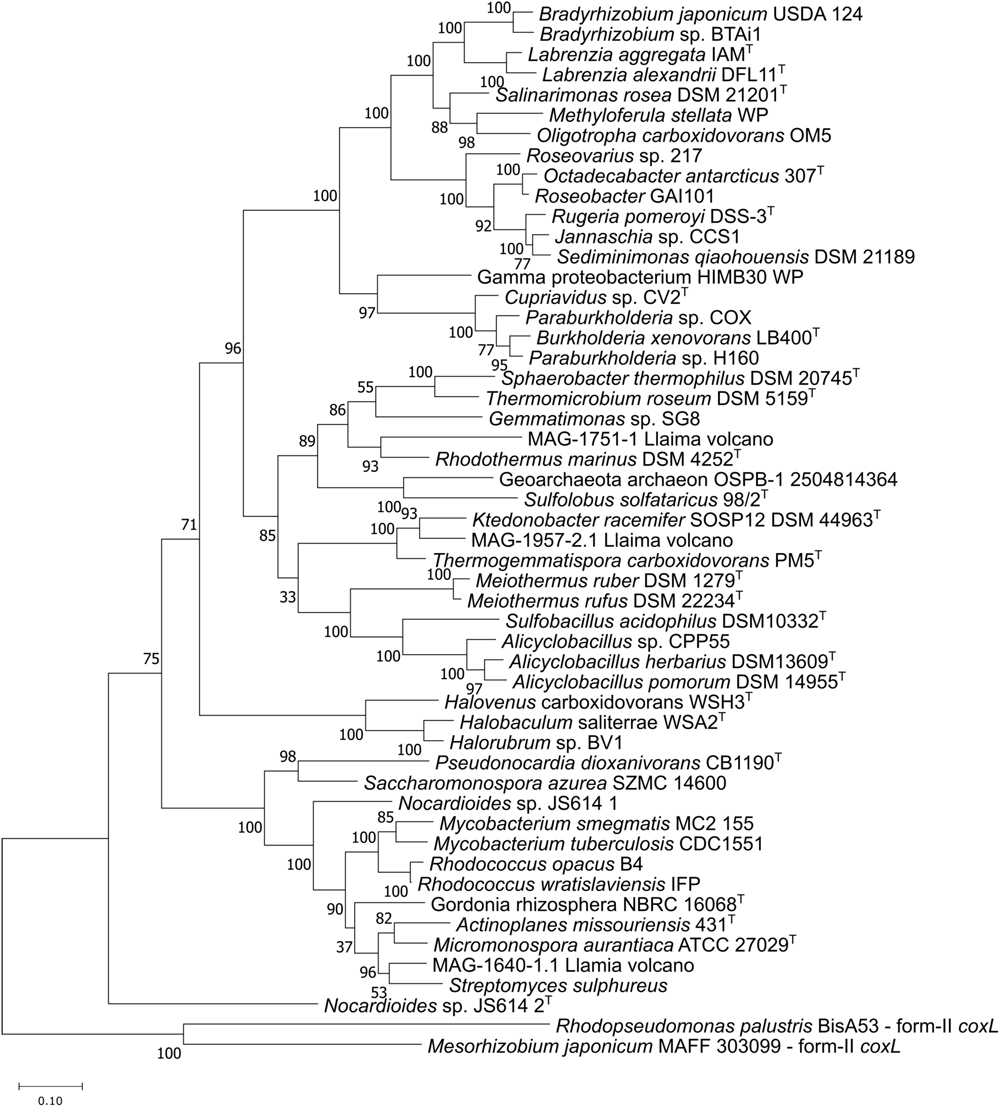
Evolutionary relatedness of translated CoxL amino acid sequences. The tree was drawn using the Maximum Likelihood method with 500 Bootstrap replicates in MEGA7. MAGs were retrieved from Hernández *et al*. (2020).

*Cupriavidus* sp. CV2^T^ and *Pb. terrae* COX only possess a limited selection of the accessory *cox* genes (*coxGDEF*), which were initially identified on the classic CODH-encoding gene cluster from *O. carboxidovorans* OM5 (Figure 2). Previous studies in *O. carboxidovorans* OM5 demonstrated that the accessory Cox proteins (CoxBC, CoxDEFGHIK) have roles in the maturation and anchoring of CODH (CoxMSL) (Meyer *et al*., 2000; Nam *et al*., 2003). The translated CoxGDEF homologs identified in these novel carboxydovores only had low identity (27.9-56.9%) with the corresponding translated amino acid sequences from strain OM5, indicating that these proteins may have differing roles. Two distinct genes were present in the novel strains when compared to strain OM5; *rcoM*, which encodes a putative heme-containing CO-sensing transcription factor (Kerby *et al*., 2008; Kerby and Roberts, 2012), and *mocA*, which encodes a molybdenum cofactor cytidyltransferase that facilitates the assembly of the molybopterin cytosine dinucleotide (MCD) cofactor required by CODH (Neumann *et al*., 2009) (Figure 2). The presence of all accessory *cox* genes is not universally essential to CO oxidation, as *Haliangeum ochraceum* is capable of expressing functional CODH with only *coxMSL* and a single accessory gene (Quiza *et al*., 2014; Weber and King, 2017). Further study would benefit greatly from the use of targeted gene deletions to verify the genes that are essential to CO oxidation by carboxydovores.

The genomes of *Cupriavidus* sp. CV2^T^ and *Pb. terrae* COX indicated that each bacterium lacked a complete CO_2_ fixation pathway. However, each genome encodes a phosphoenolpyruvate carboxylase, meaning that assimilation of a small amount of carbon from CO through anaplerotic CO_2_ fixation cannot be discounted (Braun *et al*., 2021). Carboxydovory was confirmed as incubation of each strain with elevated concentrations of CO, with or without organic carbon, failed to facilitate growth (Figure S4; Figures 3A, 3C). The genome of *Cupriavidus* sp. CV2^T^ revealed a capacity for trace gas utilisation beyond CO alone. A gene cluster encoding a putative Ni-Fe hydrogenase was detected, which had a high level of identity at the amino acid level with the functionally verified hydrogenase gene cluster from *Cupriavidus necator* H16 (Schwartz *et al*., 2003) and, additionally, shared the same genetic organisation (Figure S5). The ability to gain energy from the oxidation of both CO and H_2_ may provide a competitive advantage to carboxydovores such as *Cupriavidus* sp. CV2^T^ over other mixotrophs during the colonisation of volcanic deposits (King, 2003a). Hydrogenases are widely distributed in nature and their activity has been observed in volcanic cinders (King and King, 2012). Hydrogenases also support microbial productivity when organic carbon is limited (Greening *et al*., 2016; Islam *et al*., 2020). *Pb. terrae* COX lacked any recognisable hydrogenase-encoding genes on its genome, suggesting that it employed other methods of surviving in nutrient-poor unvegetated volcanic deposits. Evidence from genomes and studies with cultured isolates have demonstrated the use of both CO and H_2_ by many carboxydotrophs and carboxydovores (Meyer and Schlegel, 1978; Mörsdorf *et al*., 1992; King, 2003c; King and Weber, 2007; Islam *et al*., 2019; Hernández *et al*., 2020b), suggesting that the use of both trace gases may be a relatively common strategy.

### Physiological and metabolic characterisation

*Cupriavidus* sp. CV2^T^ and *Pb. terrae* COX grow using a relatively wide range of organic carbon sources (Table S2). *Cupriavidus* sp. CV2^T^ was unable to use most of the sugars tested in this study, with only weak growth observed with arabinose. Similarly, *Cupriavidus basilensis* DSM 18853 was unable to utilise glucose (Steinle *et al*., 1998). *Pb. terrae* COX, however, grew on a range of organic compounds. The growth substrate ranges observed in these novel strains may reflect their native environments. *Pb. terrae* COX was isolated from a recent volcanic deposit (lava 2015), lacking in soil or organic matter, possibly explaining the metabolic versatility of this strain (Table S2). *Cupriavidus* sp. CV2^T^, however, was isolated from soil from the 1917 volcanic stratum. This may allow the resident microorganisms to streamline their metabolism to make use of a smaller substrate range. With this in mind, it is curious to note that *Cupriavidus* sp. CV2^T^ still has a 10 Mbp genome containing many genes with predicted roles in metabolic pathways (Table S1). Although the genus *Cupriavidus* can use a wide range of substrates, including sugars in some cases (Brigham *et al*., 2010; Park *et al*., 2011; Pearcy *et al*., 2022), *Cupriavidus* sp. CV2^T^ may be unable to use sugars such as glucose due to the lack of a gene encoding phosphofructokinase, as determined by KEGG analysis on the MicroScope annotation platform.

*Cupriavidus* sp. CV2^T^ grew on 5 mM pyruvate between pH 5.0-8.0 (Figure S6), and *Pb. terrae* COX grew between pH 5.0-7.0 (Figure S7). A relatively broad pH tolerance may be a beneficial trait to volcanic bacteria in general, as volcanic ash can significantly contribute to soil acidification (Lubis *et al*., 2021). After 24 hours, *Cupriavidus* sp. CV2^T^ had grown to a statistically similar OD_600_ at pH 6.0, 7.0 and 8.0, with a small but significant decrease in growth at pH 5.0 when compared to the latter three conditions (p≤0.05) (Figure S6). No statistically significant difference was detected after 48 hours of growth (p>0.05). Additionally, no growth was observed at pH 4.0 over the course of the experiment. *Pb. terrae* COX grew to a significantly higher final OD_600_ at pH 7.0 than at any other pH (p≤0.01), and remained significantly higher over the course of the experiment (Figure S7). No statistically significant difference in growth was observed between pH 5.0 or 6.0 over the course of the experiment, and no growth was observed at pH 4.0 or 8.0. These data indicated that *Cupriavidus* sp. CV2^T^ has a broader pH tolerance than *Pb. terrae* COX, possibly making it more suited to surviving in more changeable environments, but both strains are unable to grow under more acidic conditions.

### CO consumption and tolerance of carboxydovores

Both *Cupriavidus* sp. T CV2 and *Pb. terrae* COX consumed CO when grown in liquid culture with 5 mM pyruvate (Figure 4A-4D). A significant difference in culture density for either strain during incubations 0-100,000 ppmv CO was only detected immediately after inoculation (ANOVA, *p*≤0.01), with no significant difference in growth detected at any other timepoint. These data strongly supported the use of carboxydovory by *Pb. terrae* COX and *Cupriavidus* sp. CV2^T^, as these strains were likely to be acquiring energy for maintenance from CO oxidation (King and Weber, 2007). 200 ppm CO was consumed below detection by *Pb. terrae* COX after 7 days (Figure 5), with an initial lag period of approximately 2 days before CO was consumed. Similarly, *Cupriavidus* sp. CV2^T^ did not consume CO for the first 72 hours of incubation, with 200 ppm CO consumed below detection after 7 days (Figure 4C). For each strain, CO consumption did not occur while growth on pyruvate was occurring (Figures 4 and 5), consistent with previous reports that carboxydovores switch to trace gas oxidation during stationary phase (Islam *et al*., 2019). It was previously reported that carboxydovores were unable to grow using CO as the sole source of carbon and energy due to inhibition of CO oxidation at high concentrations of CO (Hardy and King, 2001; Weber and King, 2007). For example, CO uptake by *S. aggregata* was inhibited by concentrations of CO above 1000 ppmv (Weber and King, 2007). *S. aggregata* is unlikely to encounter high concentrations of CO *in situ* as 0.1 Tg C CO yr^-1^ is actually released to the top 10 m of the ocean (Conte *et al*., 2019). The thermophilic carboxydovore *Thermomicrobium roseum* was able to oxidise CO completely at 50,000 ppmv and still exhibited the ability to consume CO at 200,000 ppmv, although it could not oxidise all of the CO (Wu *et al*., 2009). *Cupriavidus* sp. CV2^T^ and *Pb. terrae* COX were isolated at an enrichment concentration of ∼100 ppmv, lower than the affinities/requirements for carboxydotrophs (Cypionka and Meyer, 1982; Mörsdorf *et al*., 1992; King and Weber, 2007), but the tolerance of these strains for higher concentrations of CO was unknown. Geothermal activity may cause transient changes in CO concentration in volcanic ecosystems, particularly as CO production by soils appears to increase with rising temperature (Inman *et al*., 1971; Conrad and Seiler, 1980), but much of the CO available in established soils would be provided by plant roots (King and Crosby, 2002) and abiotic degradation of biomass (Conrad and Seiler, 1980). As transient increases in CO concentrations may occur in volcanic (and other geothermal) environments, the question of how *Cupriavidus* sp. CV2^T^ and *Pb. terrae* COX would respond to elevated CO was investigated.

**Figure 4.**
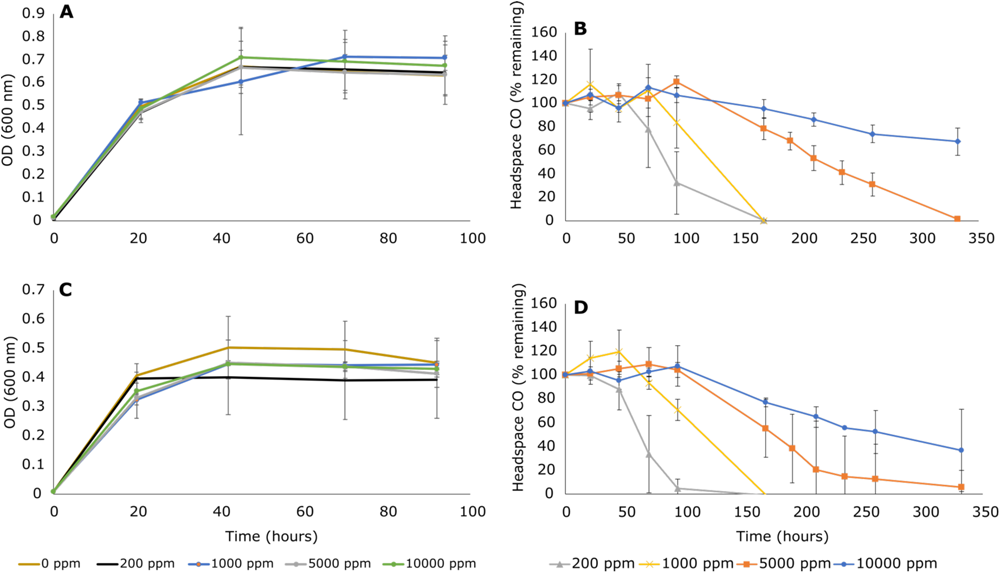
Growth of, and CO consumption by, bacterial strains. When determining statistically significant consumption of CO during CO tolerance experiments, one-way ANOVA was used to compare all timepoints within a given concentration. A) Growth of *Cupriavidus* sp. CV2^T^ with 5 mM pyruvate combined with varying concentrations of headspace CO (0-100,000 ppmv) (OD_600_). B) Headspace CO concentrations (ppm) during growth with 5 mM pyruvate by *Cupriavidus* sp. CV2^T^, Y-axis values are presented on a Log_10_ scale. C) Growth of *Pb. terrae* COX with 5 mM pyruvate combined with varying concentrations of headspace CO (0-100,000 ppmv) (OD_600_). D) Headspace CO concentrations (ppmv) during growth with 5 mM pyruvate by *Pb. terrae* COX, Y-axis values are presented on a Log_10_ scale. Bars represent mean values with standard deviation of independent triplicate incubations for each substrate.

**Figure 5.**
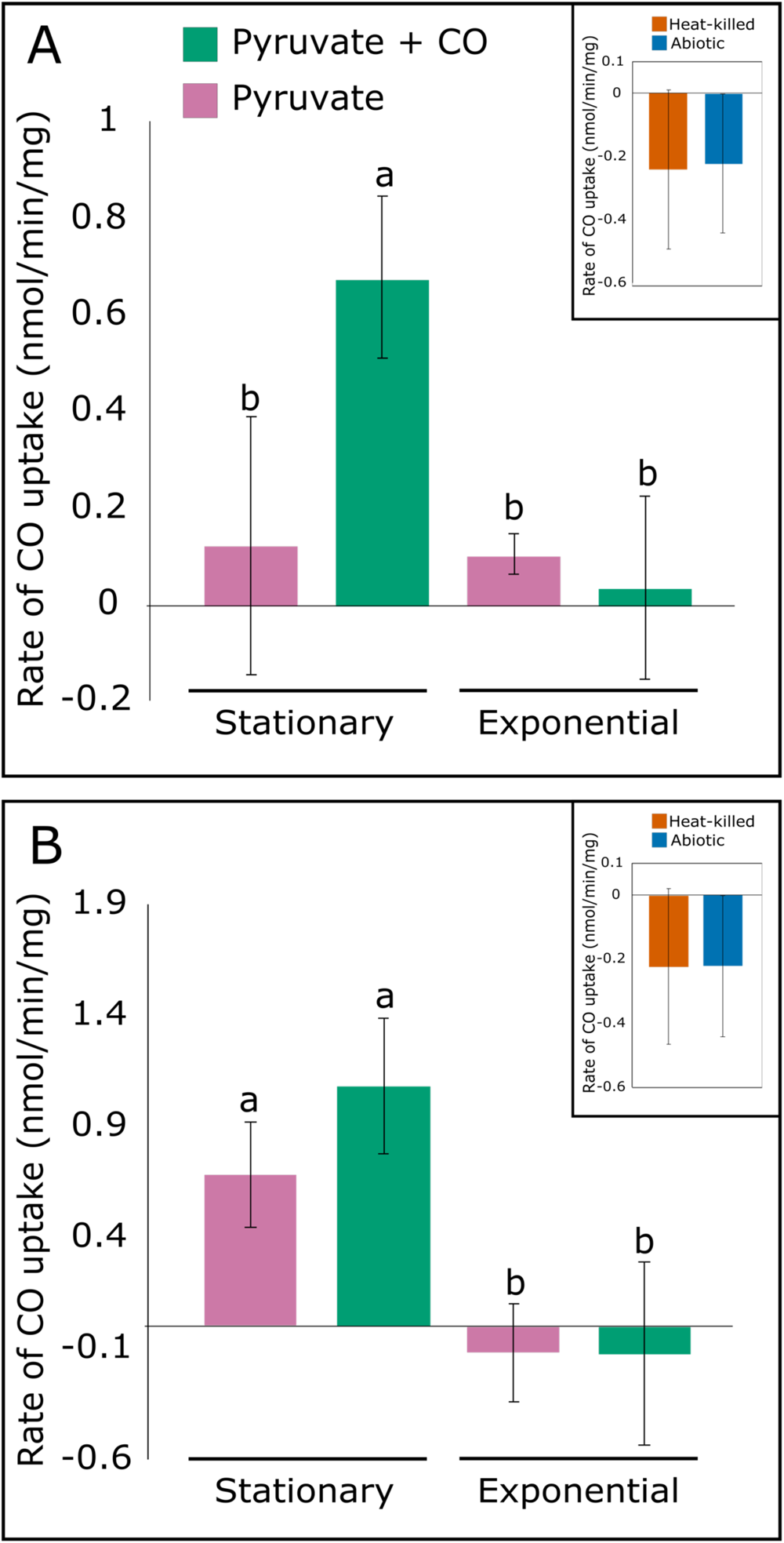
Rates of CO uptake by (A) *Cupriavidus* sp. CV2^T^ and (B) *Pb. terrae* COX at stationary phase or exponential phase, varied according to the growth substrate. Bars represent mean values with standard deviation of independent triplicate incubations for each substrate. Different letters above the bars indicate significant differences determined by one-way ANOVA followed by a Tukey post hoc test.

CO uptake was observed between 100-10,000 ppmv CO (Figure 4), but not at 100,000 ppmv CO (data not shown), and notable differences in the lag period before CO consumption began were observed for each strain at different concentrations of CO (0-10,000 ppmv). *Cupriavidus* sp. T CV2 consumed a significant amount of CO from the headspace of vials with 200 ppmv CO after 94 hours (one-way ANOVA: p≤0.001), 168 hours for 1,000 ppmv CO (p≤0.001), and 210 hours for 5,000 ppmv CO (p=5.68×10^-9^) and 10,000 ppmv CO (1% v/v) (p=3.32×10^-5^). *Pb. terrae* COX consumed a significant quantity of headspace CO after 70 hours with an initial concentration of 200 ppmv (one-way ANOVA: p≤0.001), 168 hours for 1000 ppmv (p≤0.01), 190 hours for 5,000 ppmv (p=2.47×10^-7^) and did not consume a significant quantity of CO from the headspace with 10,000 ppmv (1% v/v) CO (p>0.05). The increasing lag before CO consumption with increasing initial CO concentrations, without any significant detriment to growth that could indicate broader toxicity to cellular processes (Figures 4A, 4C), indicated that elevated CO concentrations are inhibitory to the activity of CODH in *Cupriavidus* sp. CV2^T^ and *Pb. terrae* COX (Figures 4B, 4D).

*Cupriavidus* sp. CV2^T^ appears to have a greater tolerance for elevated CO than *Pb. terrae* COX, as *Cupriavidus* sp. CV2^T^ exhibited a shorter lag period before consuming a significant quantity of CO at 5,000-10,000 ppmv CO (Figures 4B, 4D), and *Pb. terrae* COX did not consume any CO from the 100,000 ppmv headspace during the test period. *Pb. terrae* COX exhibited the shortest lag period before consuming a significant quantity of CO at 200 ppmv initial concentration, indicating that this strain may be more effective at utilising smaller concentrations of CO. The levels of CO experienced by *Cupriavidus* sp. CV2 and *Pb. terrae* COX *in situ* are unknown. However, these data can be justified when considering the environmental differences between the sites of isolation for each strain, as the lack of soil in the deposit from 2015 may have required *Pb. terrae* COX to rely more heavily on reduced gases such as CO for persistence. *Cupriavidus* sp. CV2^T^ and *Pb. terrae* COX have a similar tolerance for CO when compared to carboxydovores such as *T. roseum* (Wu *et al*., 2009), and a greater tolerance than marine CO degraders such as *S. aggregata* (Weber and King, 2007). The autotrophic non-CO degrader *Cupriavidus necator* H16 was able to tolerate 50% (v/v) CO following laboratory evolution, but the wild-type strain was inhibited by ≥15% (v/v) CO (Wickham-Smith *et al*., 2023). As CO inhibited autotrophic growth in *C. necator* H16 rather than CODH activity, it is unknown whether tolerance to elevated CO could occur in a similar laboratory evolution study with *Cupriavidus* sp. T CV2. Little information is available in the literature to indicate the typical level of tolerance to CO in *Paraburkholderia* spp. Overall, these data demonstrate that *Cupriavidus* sp. CV2 and *Pb. terrae* COX exhibit lower CO tolerance compared to many carboxydotrophs (Cypionka and Meyer, 1982; Mörsdorf *et al*., 1992). However, their tolerance levels are still higher than what appears necessary when considering ambient CO concentrations ranging from 60-300 ppbv (King, 2003b), even with transient increases in CO due to abiotic processes. Further studies are required to better understand whether CO tolerance in these strains depends on CODH activity, whether these strains benefit from CO-insensitive *o-*type cytochromes (Meyer *et al*., 1986), and whether CO tolerance is influenced by the ecology of CO-degrading bacteria.

### Regulation of CO consumption by growth conditions and growth phase

The lack of CO consumption during the initial period of growth on pyruvate (Figure 4A-4D) indicated that each carboxydovore switched from heterotrophic growth on organic compounds (pyruvate) to chemolithoheterotrophic energy acquisition through CO oxidation when organic carbon became limited. This would be consistent with previous observations by Islam *et al*. (2019) who showed that the thermophilic carboxydovore *T. roseum* underwent extensive metabolic remodelling during the shift to stationary phase, favouring processes such as H_2_ and CO oxidation. Mixotrophic use of CO during heterotrophic growth has also been reported in carboxydotrophs (Kiessling and Meyer, 1982; Kim and Kim, 1989), confirming that such metabolism is not limited to the carboxydovores. Additionally, CO-induced *cox* gene transcription by the carboxydotroph *O. carboxidovorans* OM5 occurred specifically in the absence of organic substrates (Santiago *et al*., 1999). The conditions under which *Cupriavidus* sp. CV2^T^ and *Pb. terrae* COX consume CO were investigated by growing each strain with either pyruvate or a combination of pyruvate and CO, then testing the rate of CO uptake during exponential or stationary phase (Figure 5A, 5B). The rate of CO oxidation increased significantly during stationary phase compared to exponential phase for cells that were grown with a combination of pyruvate and CO (one-way ANOVA: p≤0.01). Little to no CO uptake was detected in cells harvested at exponential phase for either strain, consistent with the use of CO oxidation as a starvation response by carboxydovores. A key difference was seen between *Cupriavidus* sp. CV2^T^ and *Pb. terrae* COX during the shift to stationary phase. *Cupriavidus* sp. CV2^T^ oxidised very little CO during stationary phase after growth with pyruvate alone (Figure 5A), likely indicating that CODH expression requires the presence of CO. *Pb. terrae* COX, however, oxidised CO at a very similar rate during stationary phase regardless of the growth condition (one-way ANOVA p≥0.05) (Figure 5B), indicating that CO uptake by *Pb. terrae* COX may not be regulated by the presence of CO but, rather, by the presence or absence of other organic carbon sources.

The regulation of CODH expression is highly variable between different carboxydotrophs and carboxydovores. CODH expression is constitutive in some carboxydotrophs such as *Pseudomonas carboxydoflava* and *Hydrogenophaga pseudoflava* (Kiessling and Meyer, 1982; Meyer *et al*., 1986; Kang and Kim, 1999), although *P. carboxydoflava* begins to use gases such as CO and H_2_ only as supplementary energy sources (mixotrophy) when supplied with heterotrophic substrates. This was suggested to support *P. carboxydoflava* in assimilating surplus organic carbon (Kiessling and Meyer, 1982). Conversely, other carboxydotrophs have demonstrated inducible expression of CODH (Kiessling and Meyer, 1982; Santiago *et al*., 1999). Repression of CODH activity by organic carbon sources was previously demonstrated, as CO uptake in the carboxydovore *S. aggregata* and carboxydotroph *O. carboxidovorans* was reduced or eliminated (Santiago *et al*., 1999; Weber and King, 2007). Weber and King (2007) suggested that glucose (or glucose metabolites) exerted allosteric repression on CODH expression in *S. aggregata,* even though this bacterium could not grow on glucose. Carboxydovores such as *T. roseum* and *Mycobacterium smegmatis* switch to CO oxidation during starvation (Cordero *et al*., 2019; Islam *et al*., 2019), suggesting that the regulation of CODH expression is dependent on levels of organic carbon. Novel carboxydovores *Cupriavidus* sp. CV2^T^ and *Pb. terrae* COX both demonstrate inducible expression of CODH, as activity was only observed during stationary phase (Figures 5A, 5B). The difference in CO uptake between growth substrates for each strain is curious. Consistent with previous observations that carboxydovores derive supplementary energy from CO oxidation during nutrient limitation (Cordero *et al*., 2019; Islam *et al*., 2019), *Pb. terrae* COX, itself a resident of a nutrient-poor unvegetated volcanic deposit, appeared to have developed a general CO scavenging response during the shift to stationary phase (Figure 5B). *Cupriavidus* sp. CV2^T^, isolated from a 1917 volcanic deposit with higher organic matter abundance, seemed to adopt a more conservative approach, expressing CODH only during the stationary phase if CO was present (Figure 5A). Detailed analysis of the metabolism and physiology of other carboxydovores may provide greater insight into the mechanisms of control of CO oxidation.

### Conclusions

This study demonstrates an effective strategy for enriching and isolating carboxydovore bacteria from volcanic soils. In this work two new strains are presented; *Cupriavidus* sp. CV2^T^ and *Pb. terrae* COX. These strains consumed CO specifically during stationary phase, with *Cupriavidus* sp. CV2^T^ employing a more conservative regulation strategy where CODH expression required the presence of CO, while *Pb. terrae* COX expressed CODH at stationary phase regardless of the presence of CO. Putative transcriptional regulators were identified with the *coxMSL* gene clusters, which could be studied with the aim of better understanding the environmental triggers of CO uptake. The tolerance of these strains to elevated CO concentrations is similar to other carboxydovores from geothermal environments, and their metabolic versatility and pH tolerance likely contribute to the successful colonisation of hostile volcanic sediments.

Further genomic comparisons of the isolate CV2^T^ using MIGA, dDDH, ANI, and a set of 77 house-keeping genes using autoMLST, revealed similarities below the recommended ANI species-level thresholds (95%, Richter and Rosselló-Móra, 2009) and DDH values (70%, Tindall *et al*., 2010) when the type strain genome sequences were used as query. *Cupriavidus* sp. CV2^T^ is the representative of a novel species of the genus *Cupriavidus* within the order *Burkholderiales* (Phylum *Pseudomonadota*). Based on this characterisation, we propose *Cupriavidus* sp. CV2^T^ as a new species, *Cupriavidus ulmosensis* sp. nov.

### Description of Cupriavidus ulmosensis sp. nov

*Cupriavidus ulmosensis* sp. nov. (ul.mo.sen’sis. N. L. masc n. *ulmosensis*, of the Parque Valle Los Ulmos, referring to Parque Valle Los Ulmos in Chile, park named after the ulmo tree in Calbuco volcano, where the strain was isolated) The type strain *Cupriavidus ulmosensis* CV2^T^ (= NCIMB 15506^T^, = CECT 30956^T^), was isolated from soil of Calbuco volcano in the Los Lagos Region, Chile. The genome is characterized by a size of 10.31 Mb and has a G + C content of 64.8 mol%. Gram-negative, rod-shaped aerobic heterotrophic bacterium. The isolate contains CODH-related genes and can grow in the presence of CO (200 – 100,000 ppmv CO). Cells grow at 30°C, at pH 4.0-8.0 (optimally at pH 8) and with 5 mM of pyruvate in VL55 medium (pH 5.5).

## MATERIALS AND METHODS

### Targeted enrichment and isolation of carboxydovores from volcanic soil

Soil samples were collected in February 2022 from Calbuco volcano (41.3304° S, 72.6087° W), Chile. Samples were taken from a lava stratification (Fig S1) formed by eruptions in 1893, 1917, 1961 and 2015 (the most recent eruption), characterised by different levels of soil development. Physical details of the lava stratification have been reported previously (Romero *et al*., 2021). Soil samples (*in situ* pH ∼5.5) were collected in triplicate at a 30 cm horizontally into the vertical surface of the lava strata except for the topmost layer, which had only volcanic rocks. Grasses and weeds (Figure S1) were removed before sampling. In this recent site (2015, approximately 5 cm of volcanic rocks were removed before sampling to a depth that did not cross the horizon of the 1961 stratum. Samples were stored in polyethylene bags (ziplok bags) until their transport to the laboratory in Chile, where they were refrigerated at 4 °C before shipping them to our laboratory in the UK. The carboxydovores discussed in this study were isolated from samples after the lava event in 2015 and 1917.

Slurries were prepared by mixing volcanic soils/sediments with VL55 medium (DSMZ recipe 1266) at a ratio of 1g:1ml, supplemented with 1 µl ml^-1^ vitamin solution (DSMZ recipe 1266). Carbon sources were ∼100 ppmv CO and 0.5 mM pyruvate to support mixotrophy (supplied as sodium pyruvate). The slurries were incubated at 30 °C with shaking at 100 rpm. Headspace CO was measured daily as described above until it became undetectable, then the headspace was amended with approximately 100 ppmv CO. Following the consumption of ∼300 ppm CO in total, 1 ml of the enrichment medium (without soil matter) was transferred to 11 ml of fresh VL55 with 0.5 mM pyruvate and 100 ppmv CO. Headspace analysis and amendments with CO were conducted as described above until a further 300 ppm CO was consumed. The enrichment medium was diluted across a 10-fold series to 10^-5^ and spread on VL55 plates, solidified with 1.5% (w/v) Bacto agar (Fisher Scientific) and amended with 0.5 mM pyruvate.

Colonies were screened for the presence of form-I *coxL*, the active site-containing component of CODH, using primers OMPf (5’-GGCGGCTT[C/T]GG[C/G]AA[C/G]AAGGT-3’) and O/BR (5’- [C/T]TCGA[T/C]GATCATCGG[A /G]TTGA-3’) reported by King (2003). The PCR protocol was conducted as described previously (King, 2003b) with modifications to use colony biomass as the template. 5% (w/v) DMSO and 0.23% (w/v) BSA were added to the DreamTaq PCR mastermix (ThermoScientific, Waltham, MA) and the initial denaturation was maintained at 94 °C for 10 minutes to lyse cells. Colonies that contained form-I *coxL* were re-inoculated in sterile VL55 medium with 2 mM pyruvate and 100 ppmv CO and incubated at 30 °C for up to 2 weeks with shaking at 100 rpm. Cultures that consumed CO were serially diluted across a 10-fold series to 10^-5^ and spread on solid VL55 agar with 2 mM pyruvate. Colony morphology was examined for evidence of contamination and colonies were re-tested for the presence of form-I *coxL*. This process was repeated until axenic cultures of CO degrading bacteria were obtained. Two isolates were further characterised in this study (see below) and named *Cupriavidus* sp. CV2^T^ and *Paraburkholderia terrae* COX. *Cupriavidus* sp. CV2^T^ and *Pb. terrae* COX were cultivated in VL55 medium (pH 5.5, matching the pH of the original soils). Cells were grown at 30 °C in 120 ml vials using 20 ml medium with shaking at 150 rpm. CO consumption was measured by gas chromatography.

### Extraction of DNA from volcanic soils and sediments

Total DNA from the soil samples was extracted using a DNeasy PowerSoil Pro kit (Qiagen) according to the manufacturer’s instructions. DNA was amplified with primers 341F (5’- CCTAVGGGRBCCASCAG-3’) and 806R (5’-GGACTACNNGGGTATCTAAT-3’) and 16S rRNA gene sequencing was done using Illumina PE250 at Novogene, UK. The identification of Amplicon Sequence Variants (ASV), removal of chimeras and taxonomic assignment using the RDP classifier (threshold 80%) were carried out with the DADA2 algorithm (Callahan *et al*., 2016) using Lotus2 software package (version 2.23, Özkurt *et al*., 2022) on Galaxy Europe (https://usegalaxy.eu).

### Cultivation of aerobic carboxydovores

The growth substrate ranges of each strain were tested by adding 5 mM of a defined carbon source to VL55 medium as the sole source of carbon and energy. Carbon sources were prepared as 1M stock solutions in distilled water and sterilised by passage through 0.2 µm syringe filters. Culture densities were recorded at 600 nm using a UV-1800 spectrophotometer (Shimadzu, UK) using two biological replicates. The pH range that supported VL55 medium was buffered to pH 4.0 using citric acid-Na_2_HPO_4_ buffer (0.1 M and 0.2 M, respectively), pH 5.0-6.0 using MES (2-(N-morpholino)ethanesulfonic acid), and pH 7.0- 8.0 using Trizima base. pH was adjusted using HCl or NaOH. To confirm that our isolates are carboxydovores (i.e. unable to grow using CO as the sole source of carbon and energy), each strain was initially grown using 5 mM pyruvate and 100 ppmv CO to induce CODH expression (see below). CO was added to the headspace of 120 ml vials, sealed using butyl rubber stoppers and aluminium crimp caps, from a 5000 ppmv stock prepared in N_2_. Cultures were sub-cultured into fresh VL55 medium with 1% (v/v) or 10% (v/v) CO as the sole carbon source, using three biological replicates per condition. These cultures were incubated for 17 days.

### Measurement of headspace CO

CO (CK Isotopes, UK) was measured by injecting 100 µl of headspace gas into an Agilent 7890A gas chromatograph fitted with an Agilent HP-Molsieve PLOT (Porous Layer Open Tubular) column (30 m length, 0.53 mm bore, 25 µm film, 7 inch cage,) at an initial oven temperature of 50 °C, programmed at a rate of 10 °C/min to 100 °C with no initial hold time, injector at 250 °C (1:2 split ratio) and flame ionization detector at 300 °C (carrier gas He, 4 ml min^-1^). Headspace CO was quantified relative to standards containing a known quantity of CO (8.33 ppmv – 110,000 ppmv) prepared in N_2_.

### Tolerance of carboxydovores to elevated CO concentrations

The two isolates were inoculated in VL55 medium with 5 mM pyruvate to support growth. The headspace of 120 ml vials was amended with 200 ppmv, 1000 ppmv, 10,000 ppmv (1% v/v) or 100,000 ppmv (10% v/v) CO, using three biological replicates per condition. Headspace CO concentrations were measured by gas chromatography as described above for a maximum of 14 days, or until all detectable CO was consumed. Culture density (OD_600_) was measured using a spectrophotometer every 24 hours until growth ceased.

### Differential CO uptake activity under different growth conditions and stages of growth

The isolates were cultivated in 400 ml VL55 medium in 2 L flasks with 5 mM pyruvate ± 200 ppmv CO, using three biological replicates per condition. 10 ml aliquots of cultures were harvested at exponential phase (18-24 hours) or stationary phase (72 hours) by centrifuging at 4000 *g* for 10 minutes at 4 °C, followed by resuspension to an OD_600_ of 4.0 in 1 ml of fresh VL55 medium. Cell suspensions were kept on ice in sealed 30 ml vials during transfer to a water bath heated to 30 °C with shaking at 150 rpm. Vials were allowed to pre-warm for 3 minutes, then 200 ppmv CO was added to the headspace from a 5000 ppmv stock of CO in N_2_. After a further 1 minute, a headspace sample was measured by gas chromatography as described above. Headspace samples were measured every 7.5 minutes until 6 samples had been taken. Abiotic VL55 medium and heat-killed cell controls were tested under the same conditions.

### Statistical analysis

Statistically significant differences between conditions were compared by one-way analysis of variance (ANOVA) followed by a Tukey post hoc test on R studio v.4.1.3.

### Genome sequencing, annotation, and analysis

DNA was extracted from the bacterial strains grown in VL55 medium with 5 mM pyruvate using the Qiagen Genomic-tip 100/G DNA isolation kit and associated DNA buffers (Qiagen) according to the manufacturer’s instructions. Genomic DNA concentration was quantified using a Qubit dsDNA HS Assay Kit (Thermo Fisher Scientific).

The strains were initially identified by amplifying the 16S rRNA gene using primers 27F/1492R (Lane, 1991), and Sanger sequencing was conducted by Eurofins Genomics (UK). Subsequently, whole genome sequencing was performed by MicrobesNG (Birmingham, UK). Genomic DNA libraries were prepared using the Nextera XT Library Prep Kit (Illumina, San Diego, USA) following the manufacturer’s protocol with the following modifications: input DNA was increased 2-fold, and PCR elongation time was increased to 45 seconds. DNA quantification and library preparation were carried out on a Hamilton Microlab STAR automated liquid handling system (Hamilton Bonaduz AG, Switzerland). Libraries were sequenced on an lllumina NovaSeq 6000 (Illumina, San Diego, USA) using a 250 bp paired end protocol. Reads were adapter trimmed using Trimmomatic version 0.30 (Bolger *et al*., 2014) with a sliding window quality cutoff of Q15. De novo assembly was performed on samples using SPAdes version 3.7 (Bankevich *et al*., 2012), and contigs were annotated using Prokka 1.11 (Seemann, 2014). The final sequences were uploaded to Microscope (https://mage.genoscope.cns.fr/microscope (accessed 05/06/2023)) for annotation (Vallenet *et al*., 2019). The form-I CODH-encoding genes (*coxMSL*) from the type strain *Oligotropha carboxidovorans* OM5 (Santiago *et al*., 1999) were used as queries in tBLASTn analysis against the newly sequenced genomes to identify putative *cox* gene clusters. The translated CoxL component was aligned against a database of corresponding CoxL sequences using MEGA7 (Kumar *et al*., 2016) to confirm the presence of the form-I CoxL active site motif (AYXCSFR) (Dunfield and King, 2004). To determine whether our two isolates were members of previously undescribed species, the genome sequence data were uploaded to the Type (Strain) Genome Server (TYGS) (https://tygs.dsmz.de) for whole genome-based taxonomic analysis. Additionally, Microbial Genomes Atlas Online (MIGA), average nucleotide identity (ANI) and average amino acid sequence identity (AAI) was calculated using the MIGA (http://microbial-genomes.org) and Enveomics (Rodriguez-R and Konstantinidis, 2016) platforms, respectively, and confirmed with the outputs from automated multi-locus species tree analysis (autoMLST) (Alanjary *et al*., 2019).

## ACKNOWLEDGEMENTS

The authors would like to thank Barbara Corrales, Pablo Saumann, and the sampling team at Parque Volcánico Valle Los Ulmos, Chile, for their assistance with sampling at Calbuco volcano and for their warm hospitality during our visit. The authors also wish to thank Professor Aharon Oren and Professor Ramon Rosselló-Móra for their valuable discussions on bacterial identification. Robin A. Dawson was supported by Royal Society Research Fellows Enhanced grants awarded to Marcela Hernández (RF\ERE\210050 and RF\ERE\231066). Marcela Hernández was supported by a Royal Society Dorothy Hodgkin Research Fellowship (DHF\R1\211076).

## SUPPLEMENTARY MATERIAL

Supplementary material for this article can be found in the online version.

## DATA AVAILABILITY STATEMENT

The genome sequences of the isolates have been deposited in the NCBI GenBank under the accession number PRJNA1001293 (SAMN36798297 for *Cupriavidus* sp. CV2^T^ and SAMN36798260 for *Paraburkholderia terrae* COX). The 16S rRNA gene Sanger sequences have also been deposited in NCBI under the accession numbers OR536588 for *Cupriavidus* sp. CV2T^T^ and OR536592 for *Pb. terrae* COX. The amplicon-sequencing data of the soils were deposited in the NCBI Sequence Read Archive (SRA) under the bioproject accession number PRJNA1036796. Both strains have been deposited in culture collections under accession numbers NCIMB 15506^T^ and CECT 30956^T^ for Cupriavidus sp. CV2^T^ and NCIMB 15505 for *Paraburkholderia terrae* COX. The CoxL database was assembled by Prof. Gary King and is available upon request.

## AUTHOR CONTRIBUTIONS

MH conceived the project and secured the funding; MH and PA collected the samples; RAD performed laboratory experiments; MH performed 16S rRNA gene amplicon sequence analysis; GK provided advice on the enrichment and isolation of bacteria; RAD and MH analysed the data; RAD and MH drafted the paper; all authors contributed to its revisions and approved the final version.

