## Supplementary Information for "Carboxydovores from the Pseudomonadota colonise volcanic soils during succession"

Running title:

CO oxidation by bacteria in volcanic soils

This supplement supplementary information includes:

Additional methods

Tables S1 and S2

Figures. S1 to S8

Table S1. Features of the genomes of *Cupriavidus* sp. CV2$^{T}$ and *Paraburkholderia terrae* COX, determined by MicroScope and RAST annotation platforms.

| **Feature** | ***Cupriavidus* sp. CV2** | ***Paraburkholderia terrae* COX** |
|---|---|---|
| Size (bp) | 10,313,057 | 10,305,210 |
| GC Content (%) | 64.8 | 62 |
| Number of Contigs | 651 | 256 |
| L50 | 52 | 62 |
| N50 | 61,988 | 101,247 |
| Completeness (CheckM - %) | 99.5 | 99.7 |
| Contamination (CheckM - %) | 2.3 | 2.4 |
| **Genes - RAST-Predicted** | | |
| No. of coding sequences (total) | 10,380 | 10,390 |
| No. of genes with assigned function | 6,563 | 6041 |
| No. of genes without assigned function | 3,817 | 4349 |
| No. of rRNAs | 4 | 8 |
| No. of tRNAs | 63 | 53 |
| No. of pseudogenes | 12 | 8 |

| **Subsystem** | ***Cupriavidus* sp. CV2** | ***Paraburkholderia terrae* COX** |
|---|---|---|
| Cofactors, Vitamins, Prosthetic Groups, Pigments | 303 | 268 |
| Cell Wall and Capsule | 39 | 46 |
| Virulence, Disease and Defense | 78 | 79 |
| Potassium metabolism | 22 | 17 |
| Phages, Prophages, Transposable elements, Plasmids | 8 | 7 |
| Membrane Transport | 137 | 98 |
| Iron acquisition and metabolism | 13 | 41 |
| RNA Metabolism | 75 | 65 |
| Nucleosides and Nucleotides | 106 | 118 |
| Protein Metabolism | 236 | 222 |
| Cell Division and Cell Cycle | 26 | 0 |
| Motility and Chemotaxis | 17 | 81 |
| Regulation and Cell signaling | 81 | 50 |
| Secondary Metabolism | 20 | 5 |
| DNA Metabolism | 119 | 101 |
| Fatty Acids, Lipids, and Isoprenoids | 279 | 216 |
| Nitrogen Metabolism | 29 | 28 |
| Dormancy and Sporulation | 1 | 1 |
| Respiration | 190 | 200 |
| Stress Response | 132 | 161 |
| Metabolism of Aromatic Compounds | 248 | 187 |
| Amino Acids and Derivatives | 769 | 622 |
| Sulfur Metabolism | 51 | 80 |
| Phosphorus Metabolism | 39 | 42 |
| Carbohydrates | 516 | 654 |

Table S2. Growth substrate range of *Cupriavidus* sp. CV2 and *P. terrae* COX. – indicates no growth; + indicates $OD_{600}$ = 0.1-0.3; ++ indicates $OD_{600}$ = 0.3-0.6; +++ indicates $OD_{600}$ = 0.6-1.0 (n=2).

| | **Level of Growth** | |
|---|---|---|
| | ***Cupriavidus* sp. CV2** | ***Paraburkholderia terrae* COX** |
| | **Sugars** | |
| Glucose | - | ++ |
| Xylose | - | +++ |
| Sucrose | - | - |
| Maltose | - | - |
| Lactose | - | + |
| L-Sorbose | - | - |
| L-Arabinose | + | ++ |
| Ribose | - | +++ |
| | **Sugar Alcohol** | |
| Sorbitol | - | ++ |
| | **Carboxylic Acids** | |
| Succinate | ++ | ++ |
| Gluconate | ++ | +++ |
| Salicylate | - | + |
| Phthalate | - | - |
| Potassium Citrate | ++ | + |
| Sodium Citrate | +++ | +++ |
| Methanesulfonate | - | - |
| Tartarate | +++ | ++ |
| Methylmalonate | - | - |
| Propionate | - | - |
| Malate | ++ | ++ |
| Glyoxylate | ++ | + |
| Formate | - | - |
| Vanillate | - | - |
| Glutamate | +++ | ++ |
| Taurine | + | + |

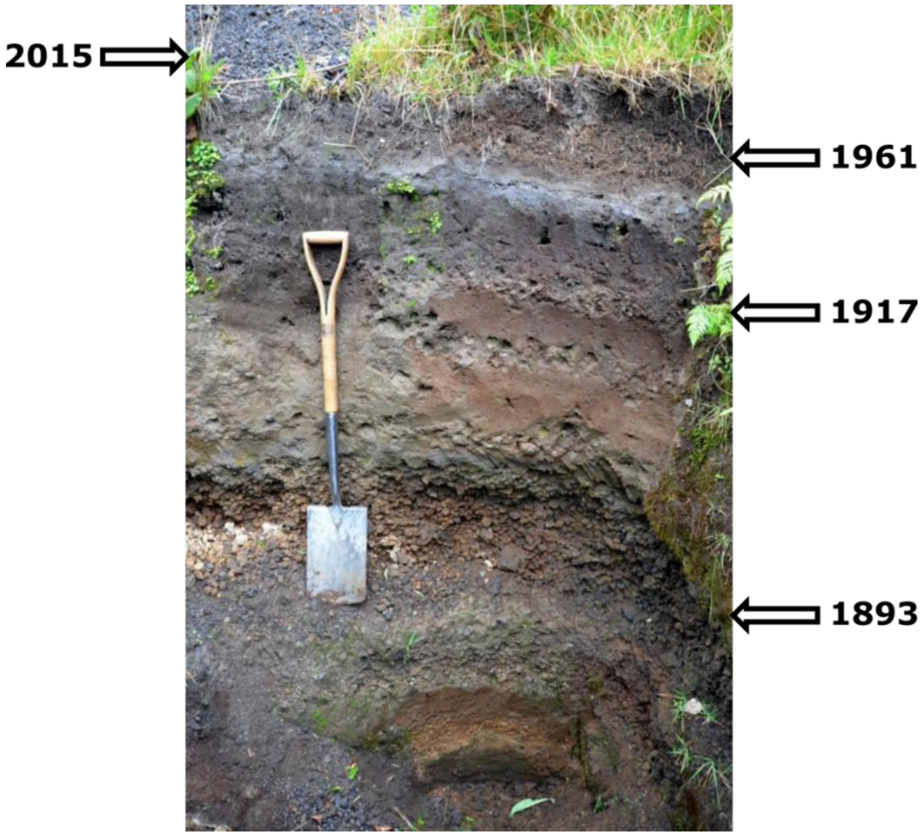


Figure S1. Calbuco volcano stratification formed by successive pyroclastic deposits. Sampling locations (indicated above) were deposited during eruptions in 2015, 1961, 1917 and 1893. Adapted from Romero *et al*. (2021).

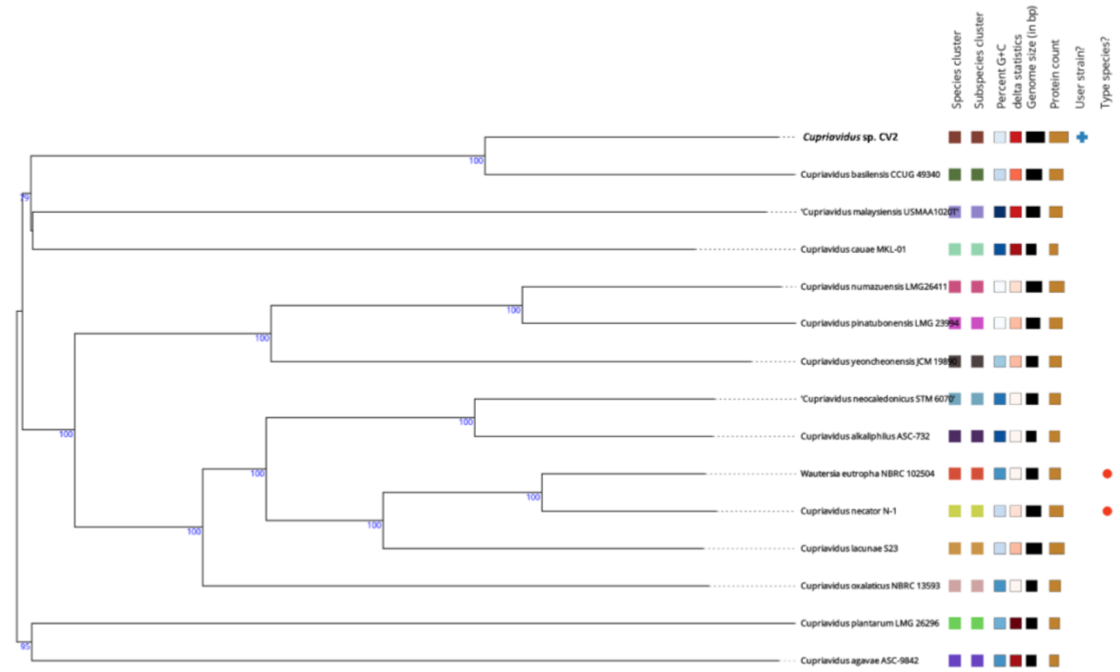


Figure S2. Whole genome-based taxonomic analysis of *Cupriavidus* sp. CV2 against representative genomes of *Cupriavidus* spp. Branch support was inferred from 100 pseudo-bootstrap replicates each.

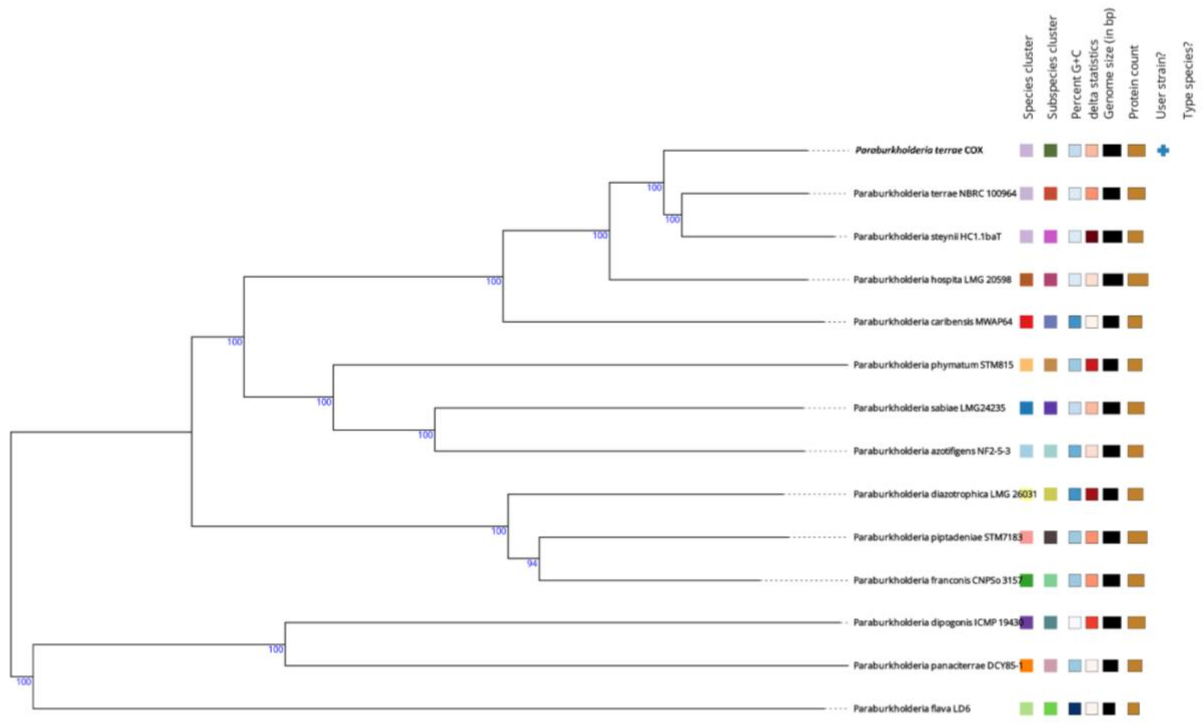


Figure S3. Whole genome-based taxonomic analysis of *Paraburkholderia terrae* COX against representative genomes of *Paraburkholderia* spp. Branch support was inferred from 100 pseudo-bootstrap replicates each.

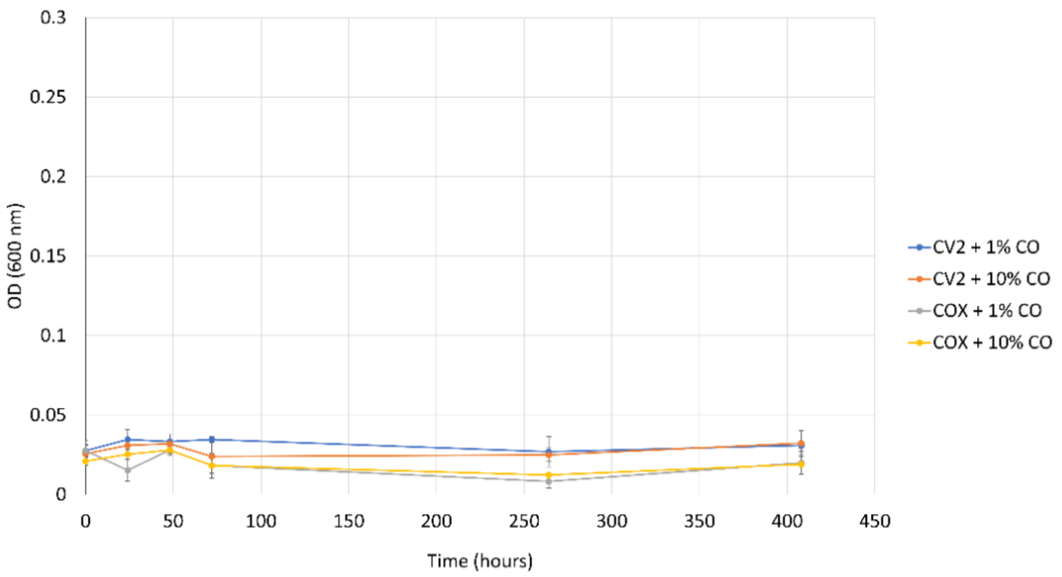


Figure S4. Confirmation of carboxydovory by incubation of *Cupriavidus* sp. CV2 and *Paraburkholderia terrae* COX with 1% (v/v) or 10% (v/v) CO as the sole source of carbon and energy (n=3). Error bars represent the standard deviation about the mean.

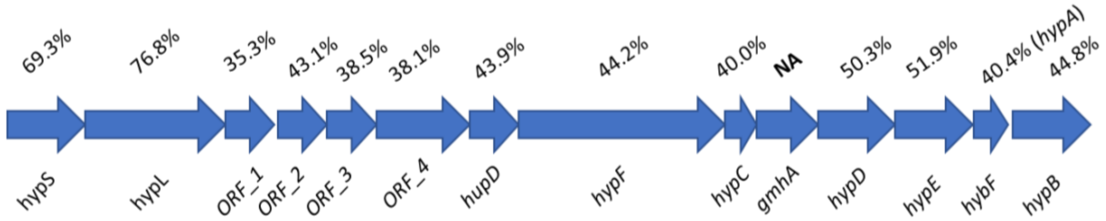


Figure S5. Gene cluster encoding a putative Ni-Fe hydrogenase in *Cupriavidus* sp. CV2. Identity of translated amino acids (%) were calculated relative to the hydrogenase gene cluster in *Cupriavidus necator* H16.

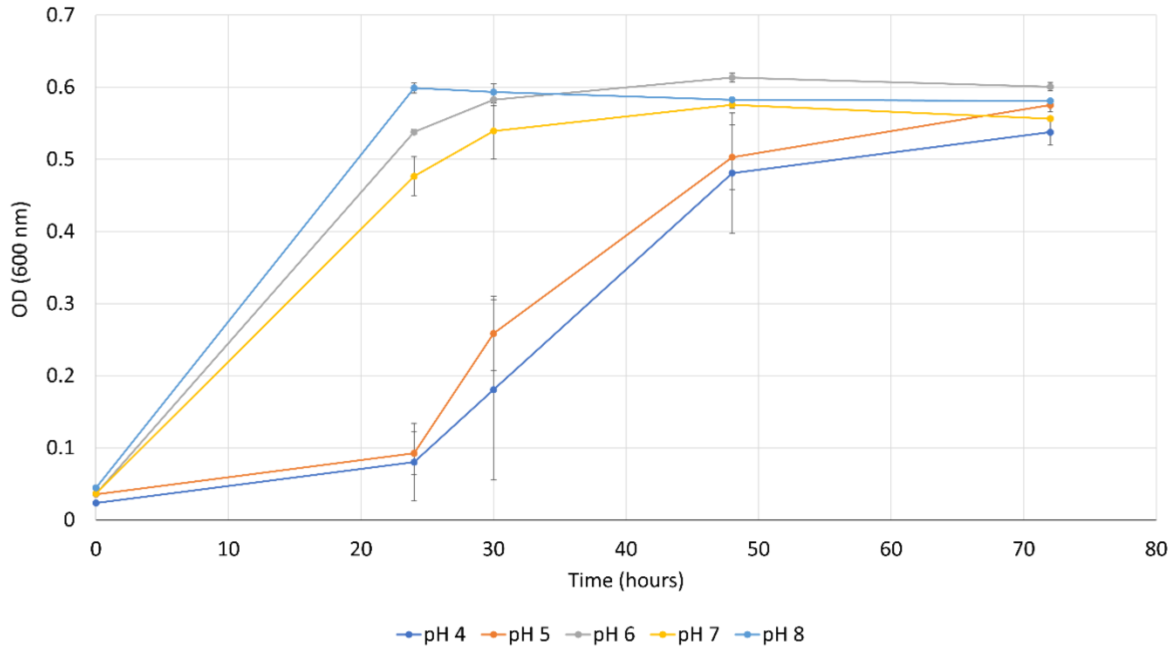


Figure S6. Growth of *Cupriavidus* sp. CV2 on 5 mM pyruvate in VL55 medium adjusted to pH 4, 5, 6, 7 or 8 using HCl and NaOH (n=3). Error bars represent the standard deviation about the mean.

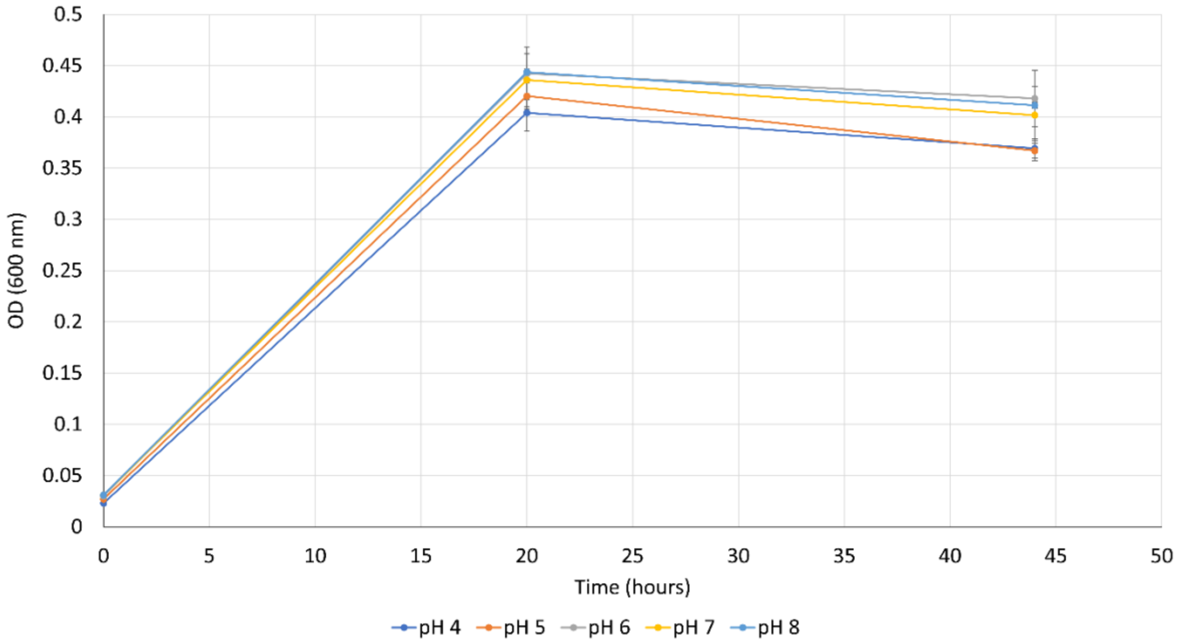


Figure S7. Growth of *P. terrae* COX on 5 mM pyruvate in VL55 medium adjusted to pH 4, 5, 6, 7 or 8 using HCl and NaOH (n=3). Error bars represent the standard deviation about the mean.
